# HIV-1 Nef Homodimerization as a Structural Mechanism for Kinase Activation and Small Molecule Inhibitor Action

**DOI:** 10.64898/2026.08.10.744017

**Authors:** Catherine E. Thomas, John J. Alvarado, Thomas E. Smithgall

## Abstract

The HIV-1 accessory protein Nef plays a central role in viral pathogenesis by enhancing viral replication, modulating cellular signaling, and evading immune recognition, making it a compelling target for therapeutic intervention. Nef lacks enzymatic activity and instead functions through diverse interactions with host cell proteins including the Src-family tyrosine kinase, Hck. Kinase activation may requires Nef homodimerization, as mutations disrupting the Nef dimer interface impair kinase activation as well as many other Nef functions. In the present study, we investigated the structural consequences of dimer interface mutations and their impact on Nef interactions with Hck regulatory domains. Using size-exclusion chromatography, multi-angle light scattering and crystallography, we found that mutations at dimer interface residues Leu112 and Phe121 abolish recombinant Nef protein dimerization while preserving the overall Nef fold, resulting in monomeric 1:1 complexes with Hck SH3 or SH3-SH2 domain proteins. These findings demonstrate that the broad phenotypic effects of interface mutations arise from loss of Nef dimerization rather than global misfolding or perturbation of SH3 binding. We also investigated the effects of small molecule Nef inhibitors on homodimer formation. These compounds, like the dimerization-defective mutations, suppress kinase activation, viral replication and restore immune recognition of HIV-infected cells. Using a SplitFAST fluorescence complementation assay, we provide direct evidence that these inhibitors disrupt Nef homodimer formation in solution. Co-crystallization of a wild-type Nef:SH3 complex with an inhibitor also prevented homodimer formation. Computational docking identified a shared pocket for six active Nef inhibitors formed by the Nef dimer interface but lost in the monomer. Together, our findings support homodimerization as a structural feature essential for many Nef functions and validate disruption of this interface as a promising therapeutic strategy against HIV-1.

## Introduction

Nef is a unique accessory protein encoded by the primate lentiviruses HIV-1, HIV-2 and SIV that plays a key role in viral pathogenesis and immune evasion (1). Lacking intrinsic enzymatic activity, Nef functions instead through diverse interactions with host cell proteins to enhance viral infectivity and replication, and to allow infected cells to avoid detection by the immune system (2, 3). Rhesus macaques infected with Nef-defective SIV display low viral loads and fail to progress to simian AIDS (4). Similarly, a subset of people living with HIV-1 that do not develop AIDS, even in the absence of antiretroviral therapy, have been shown to carry HIV-1 variants with Nef-defective alleles (4–6). These and other studies underscore the essential role for Nef in HIV/AIDS and identify this viral factor as a rational target for drug development (7). One of the most studied Nef interactions involves binding and activation of non-receptor tyrosine kinases of the Src and Tec families (8). Both kinase families share a core region consisting of the SH3, SH2 and kinase domains, with the SH3-SH2 regulatory region packing against the back of the kinase domain to control kinase activity (9). In the case of the Src-family kinase Hck, Nef-mediated kinase activation involves both SH3 domain engagement and formation of dimeric Nef:Hck complexes at the cell membrane (8). Mutations that prevent Nef-SH3 interaction as well as Nef homodimer formation disrupt kinase activation, downstream signaling, and enhancement of viral replication (10–12). However, mutations that disrupt Nef dimerization not only disrupt kinase activation but also affect several other Nef functions, including downregulation of the viral receptor CD4 and the restriction factor, SERINC5 (8, 13). These observations raise questions about broader effects of these mutations on overall Nef structure.

In the first part of this study, we investigated the impact of dimer interface mutations on the interaction and structure of Nef with the Hck SH3 and SH3-SH2 domains. Previous work has shown that engagement of Nef with either of these Hck regulatory domains results in crystal complexes with 2:2 stoichiometry in which Nef forms the dimer interface (14, 15). The Nef α-helical dimer interface in both structures involves three key residues: Leu112, Tyr115, and Phe121, with Leu112 acting as the keystone within the interface (Figure 1; residue numbering based on B-clade Nef variant, NL4-3). Here we show that recombinant Nef proteins bearing mutations at this interface still form stable complexes with the SH3 and SH3-SH2 proteins. However, size-exclusion chromatography coupled to multi-angle light scattering (SEC-MALS) analysis primarily revealed monomeric complexes with 1:1 stoichiometry, unlike the 2:2 wild-type complexes. The X-ray crystal structure of an Nef-L112A mutant core protein bound to SH3 revealed a 1:1 complex that lacked the Nef dimer interface, despite a wild-type fold of the Nef core. Taken together, these results support the conclusion that the phenotypic effects of Nef dimer interface mutations are due solely to the dimerization defect.

**Figure 1.**
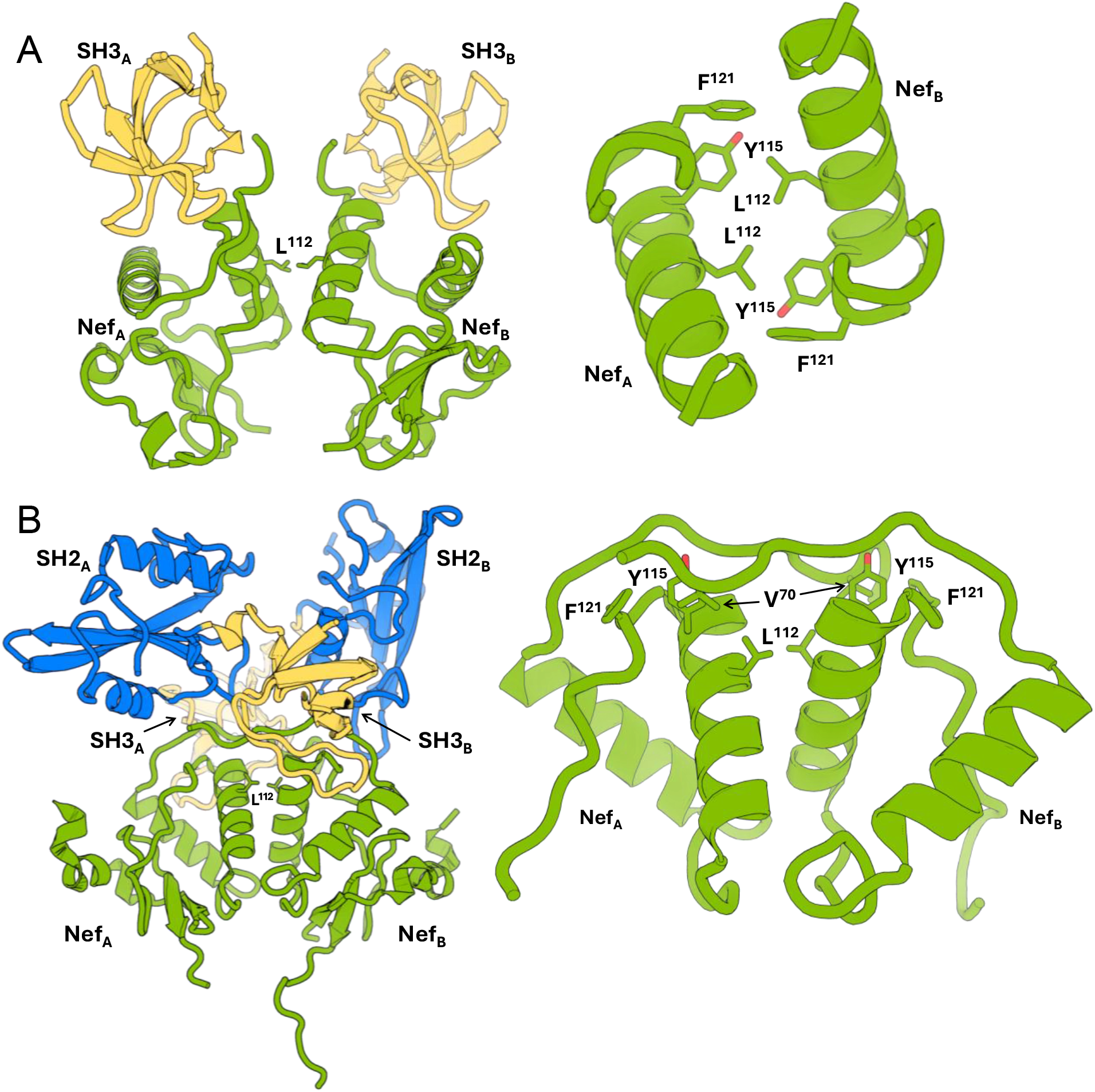
X-ray crystal structures of Nef complexed with Src-family kinase SH3 and SH3-SH2 domains. **A)** *Left*: Overall structure of Nef:SH3 complex as determined by X-ray crystallography (2.50 Å, PDB ID: 1EFN). The Fyn SH3 domain used in this structure has an Ile substitution for Arg-96 in the RT loop to model the Nef:Hck SH3 interface. Present are wild-type Nef monomers (green) and SH3 domains (yellow). The side chain of Leu112 marks the center of the helical interface. *Right*: The 1EFN Nef homodimer interface is formed by two *a*B helices and stabilized by the hydrophobic residues Leu112, Tyr115, and Phe121. **B)** *Left*: Overall structure of Nef:Hck SH3-SH2 complex as determined by X-ray crystallography (1.85 Å, PDB ID: 4U5W). Present are wild-type Nef monomers (green), SH3 domains (yellow), and SH2 domains (blue). *Right*: The 4U5W Nef homodimer interface is also formed by the two *a*B helices and stabilized by Leu112, Tyr115, and Phe121. Leu112 acts as a keystone while Phe121 and Tyr115 form a small hydrophobic pocket that engages Val70 from a loop in the opposing monomer.

The second part of our study explored the impact of small molecule inhibitors on Nef homodimer formation as well. Several classes of small molecule Nef inhibitors have been reported, the most promising of which are based on a central hydroxypyrazole core (16–18). Like the dimerization interface mutants, these compounds block multiple Nef functions, resulting in suppression of viral replication and inhibition of infectivity (16). Notably, inhibitors in this class restore MHC-I to the surface of HIV-infected primary CD4+ T cells, resulting in recognition and killing by HIV-directed cytotoxic T lymphocytes (18). These compounds also inhibit Nef-dependent activation of Hck in cell-based assays, resulting in decreased viral replication (16). Hck activation by Nef requires the homodimer (19), suggesting that the inhibitors may perturb the Nef dimer structure as well.

Using splitFAST, a reversible split fluorescent reporter protein, we developed an assay to monitor Nef homodimer formation in solution. The splitFAST system is based on a small fluorogenic protein tag (FAST) that produces a strong fluorescent signal following binding to rhodanine fluorogens (20, 21). To monitor Nef homodimer formation with this system, we expressed and purified recombinant Nef fusion proteins with non-fluorescent FAST fragments (nFAST and cFAST) fused to their C-termini. Combining the Nef-nFAST and Nef-cFAST proteins with the fluorogen resulted in complementation and fluorescence, indicative of Nef homodimer formation. Using this approach, we demonstrated that active hydroxypyrazole Nef inhibitors showed concentration-dependent reduction in fluorescence intensity, supporting disruption of Nef dimer formation. Co-crystallization of a Nef:SH3 complex in the presence of an inhibitor also prevented homodimer formation. These results provide direct evidence that this class of inhibitors suppresses multiple Nef functions by perturbing homodimer formation, disrupting a key mechanism for modulating host signaling pathways analogous to the effects of dimer interface mutations.

## Results

### Expression and purification of recombinant Nef proteins in complexes with Hck SH3 and SH3-SH2 domains

Previous studies have shown that HIV-1 Nef proteins form homodimers in the crystalline state, in solution, at model lipid bilayers, and in cell-based assays (8, 22). Mutations that disrupt the dimer interface defined by the crystal structures have broad impacts on Nef functions related to viral infectivity, replication efficiency, and immune escape (8). The pleiotropic effects of dimer interface mutations on diverse Nef functions suggest that these mutations may alter the global fold of Nef resulting in unanticipated structural changes in addition to preventing dimer formation. To address this issue, we studied the effect of Nef dimer interface mutations on complex formation with the SH3 and SH3-SH2 regulatory domains of the myeloid Src-family kinase, Hck.

Existing X-ray crystal structures of the wild-type Nef core region in complex with the SH3 domain alone or the complete SH3-SH2 regulatory region provide a structural basis for Nef homodimer formation. In the Nef:SH3 complex, which crystallizes as a 2:2 dimer, a helical dimer interface is observed that is stabilized by Nef Leu112, Tyr115 and Phe121 (Figure 1A). In the Nef:SH3-SH2 structure, which also crystallized as a 2:2 complex dimer, the same helix forms the interface with Leu112 at the center. In this structure, however, Tyr115 and Phe121 are rotated away from the interface to form a hydrophobic cup that engages Val70 in a flexible loop from the opposing Nef monomer (Figure 1B). As result, the Nef:SH3-SH2 complex is more compact with additional contacts formed between Nef and the SH2 domain as well. In both structures, the Nef interface with the SH3 domain is the same, involving the conserved polyproline helix of Nef as well Ile96 in the RT loop of the SH3 domain which contacts a hydrophobic pocket on Nef.

To explore the effects of Nef dimerization mutations on SH3 and SH3-SH2 domain interaction, wild-type and mutant Nef core domain proteins were expressed and purified in complexes with Hck SH3 and SH3-SH2 domain proteins. Three dimerization-defective Nef mutants were chosen based on their loss of kinase activation as well as impaired CD4, SERINC5 and MHC-I downregulation: L112A, L112D, and F121A (10, 12, 13, 16). Each complex was purified by affinity chromatography followed by gel filtration. SEC-MALS was then used to estimate the molecular weight and stoichiometry of each complex (Table 1). Based on the estimated molecular weights, wild-type Nef formed monodisperse complexes with SH3 or SH3-SH2 with 2:2 stoichiometry, consistent with the crystal structures (Figure 1). By contrast, the estimated molecular weights with each Nef mutant are consistent with monodisperse 1:1 monomeric complexes with SH3 and SH3-SH2. One exception was the Nef-L112A:SH3 complex. The observed molecular weight of 40 kDa corresponds more closely to that of two Nef core molecules plus one SH3 domain, a configuration previously observed in an X-ray crystal structure of the Nef core bound to the wild-type Fyn SH3 domain (PDB: 1AVZ) (23). In addition, the Nef-L112A mutant retains partial activity in some biological assays, such as downregulation of the SERINC5 restriction factor (13), suggesting some retention of homodimer formation under biological conditions. Nevertheless, all three Nef mutant complexes with SH3 displayed much smaller hydrodynamic radii than the wild-type complex. Overall, these results demonstrate that dimer interface mutations affect Nef quaternary structure without interfering with SH3 domain engagement. Note that mutagenesis of Nef Tyr115 resulted in poor yields of soluble recombinant protein, suggesting that changes to this position result in destabilization of the Nef core fold. For this reason, the Tyr115 mutant was not pursued further.

**Table 1.** SEC-MALS Analysis of wild-type and mutant Nef:Hck Complexes. SEC-MALS was performed using a Wyatt Technology instrument consisting of a DAWN multi-angle light scattering detector, a DLS-DAWN dynamic light scattering module for determination of hydrodynamic radius, and an Optilab refractometer. Purified Nef:SH3 and Nef:SH3-SH2 complexes were analyzed on a Superdex 75 Increase 10/300 GL column and data were recorded for 50 min at a flow rate of 0.5 mL/min at room temperature. Data analysis was performed with ASTRA 7.3.2 software (Wyatt Technology) which provides the estimate of the standard error.

| **Protein Complex** | **Polydispersity** | **Expected Molecular Weight (kDa)** | **Observed Molecular Weight (kDa)** | **Predicted Stoichiometry Nef:Partner** |
| --- | --- | --- | --- | --- |
| Nef-L112D:SH3 | 1.0 ± 2.6% | 26.0 | 24.5 ± 0.5% | 1:1 |
| Nef-L112A:SH3 | 1.0 ± 4.2% | 26.0 | 40.8 ± 1.2% | 2:1 |
| Nef-F121A:SH3 | 1.0 ± 2.2% | 26.0 | 29.8 ± 0.5% | 1:1 |
| Nef-WT:SH3 | 1.0 ± 10.8% | 52.0 | 44.2 ± 9.9% | 2:2 |
| Nef-L112D:SH3-SH2 | 1.0 ± 3.6% | 38.1 | 40.4 ± 1.0% | 1:1 |
| Nef-L112A:SH3-SH2 | 1.0 ± 3.6% | 38.1 | 38.9 ± 1.1% | 1:1 |
| Nef-F121A:SH3-SH2 | 1.0 ± 5.2% | 38.1 | 42.8 ± 1.9% | 1:1 |
| Nef-WT:SH3-SH2 | 1.0 ± 1.0% | 76.2 | 76.3 ± 1.8% | 2:2 |

### Loss of the Nef Homodimer Does Not Alter Nef Core Structure or SH3 Binding

To better understand the impact of dimer interface mutations on the global fold of Nef and the SH3-binding interface, we performed crystallization screens for each of the mutant Nef:SH3 and Nef:SH3-SH2 complexes shown in Table 1. Of these, the Nef-L112D:SH3 (22) and Nef-L112A:SH3 complexes yielded diffracting crystals from which we determined X-ray structures to 2.65 Å and 2.15 Å resolution, respectively (Figure 2 and Table 2). Both structures crystallized in space group P412_1_2 with the SH3 domain bound to the mutant Nef core proteins in the same manner as wild-type Nef. However, no evidence of the Nef dimer interface was observed in the density, with each complex demonstrating a 1:1 Nef to SH3 stoichiometry.

**Figure 2.**
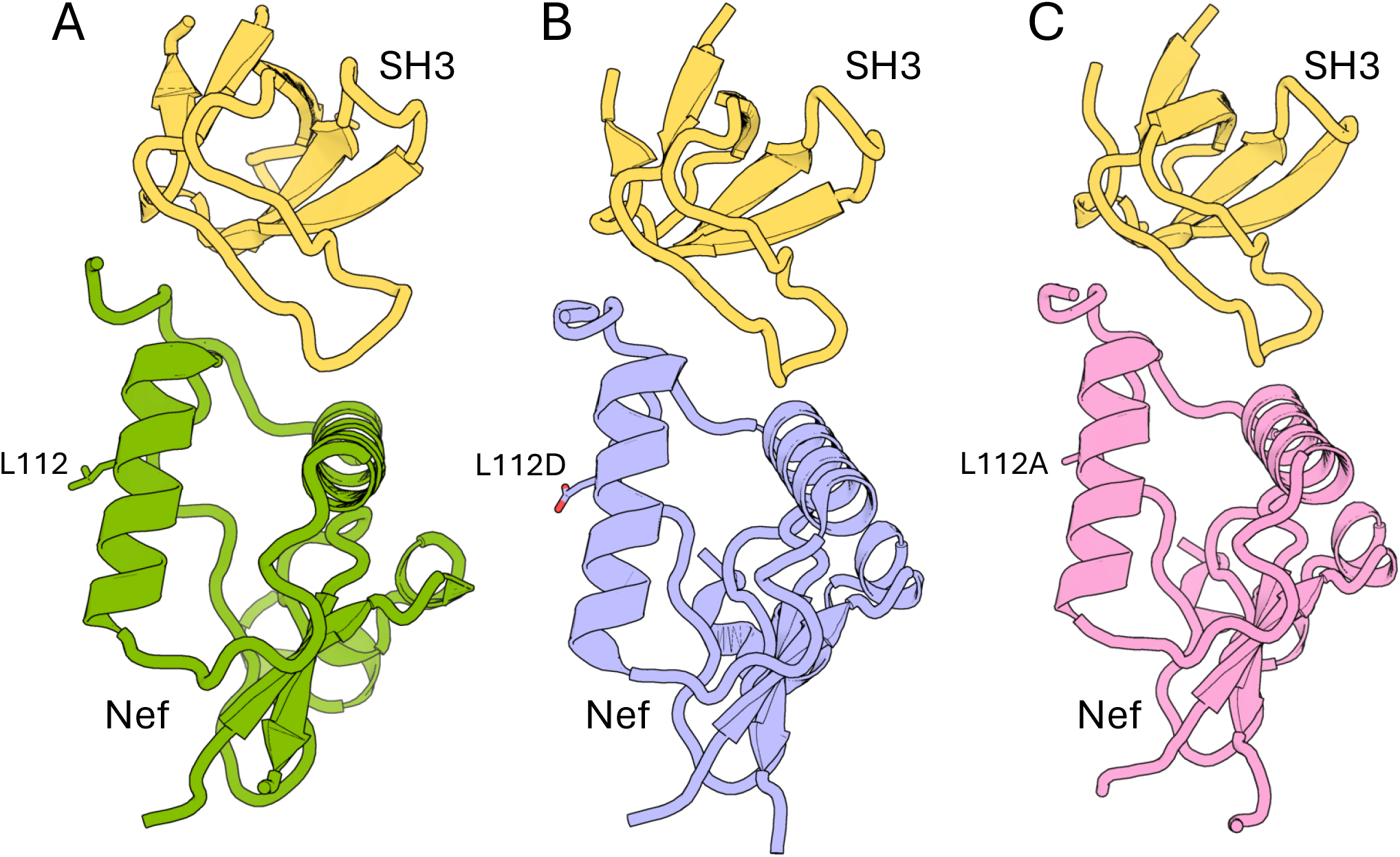
X-ray crystal structures of wild type and dimerization-defective Nef mutants in complexes with Src-family kinase SH3 domains. **A.** Overall structure of the wild-type Nef core in complex with the Fyn SH3-R96I domain (PDB ID: 1EFN). **B,C.** Overall structures of Nef-L112D (periwinkle) and Nef-L112A (pink) mutants in complex with the Hck SH3 domain (PDB IDs 8F2P and 9Y58, respectively).

**Table 2.** X-ray data collection and refinement statistics. Statistics for the highest resolution shell are shown in parentheses.

| Parameter | Nef L112A:SH3 | Nef-WT:SH3 + 7098 |
| --- | --- | --- |
| X-Ray source | 23-IDD | University of Pittsburgh |
| PDB I.D. | 9Y58 | 9Y9K |
| Resolution (Å) | 37.71-2.251 (2.33-2.25) | 36.24 – 2.34 (2.58 – 2.34) |
| Wavelength (Å) | 1.033167 | 1.541780 |
| Space group | P 4<sub>1</sub> 2<sub>1</sub> 2 | P 4<sub>1</sub> 2<sub>1</sub> 2 |
| **Cell dimensions** |  |  |
| a, b, c (Å) | 84.33, 84.33, 83.16 | 80.71, 80.71, 82.36 |
| $\alpha$, $\beta$, $\gamma$ (°) | 90, 90, 90 | 90, 90, 90 |
| Mean ($I/\sigma$) | 21.1 (0.95) | 14.62 (1.00) |
| Redundancy | 12.4 (12.8) | 13.0 (10.1) |
| Completeness (%) | 99.90 (99.74) | 98.76 (97.03) |
| $R_{\text{merge}}$ | 0.0516 (2.548) | 0.1255 (2.033) |
| CC (1/2) | 1 (0.553) | 0.999 (0.802) |
| **Refinement** |  |  |
| Resolution (Å) | 37.61 - 2.251 | 36.24 – 2.34 |
| No. of reflections | 14743 (1440) | 11843 (2845) |
| $R_{\text{work}}/R_{\text{free}}$ | 0.2381/0.2727 | 0.2409/0.2840 |
| **Number of non-H atoms** |  |  |
| Protein | 1355 | 1391 |
| Solvent | 4 | 31 |
| **Protein molecules** |  |  |
| Nef SF2core | 1 | 1 |
| Hck SH3 | 1 | 1 |
| **RMS deviations** |  |  |
| bonds length (Å) | 0.007 | 0.003 |
| Bond angles (°) | 0.84 | 0.48 |
| **Ramachandran** |  |  |
| Favored (%) | 97.45 | 96.89 |
| Allowed (%) | 2.55 | 3.11 |
| Outlier (%) | 0.00 | 0.00 |
| **Average B factor (Å<sup>2</sup>)** |  |  |
| protein | 76.66 | 65.42 |
| solvent | 67.80 | 57.63 |

Despite the loss of the Nef dimer, the overall fold of the Nef core region and the SH3 interface of both mutant structures remained the same. In the wild-type Nef:SH3 structure, the polyproline helix of Nef, flanked by Pro72 and Pro75, engages the surface grooves of the SH3 domain, packing against a series of conserved SH3 hydrophobic residues. This interaction is stabilized by a salt bridge formed between Arg77 of Nef and Asp100 in the RT loop of the SH3 domain. The RT loop also stretches across the Nef surface, with SH3 Ile96 engaging a small hydrophobic pocket formed by Phe90, Trp113 and Tyr120 of Nef (Figure 3A). These interactions are sustained in both Nef-L112D:SH3 (Figure 3B) and Nef-L112A:SH3 (Figure 3C).

**Figure 3.**
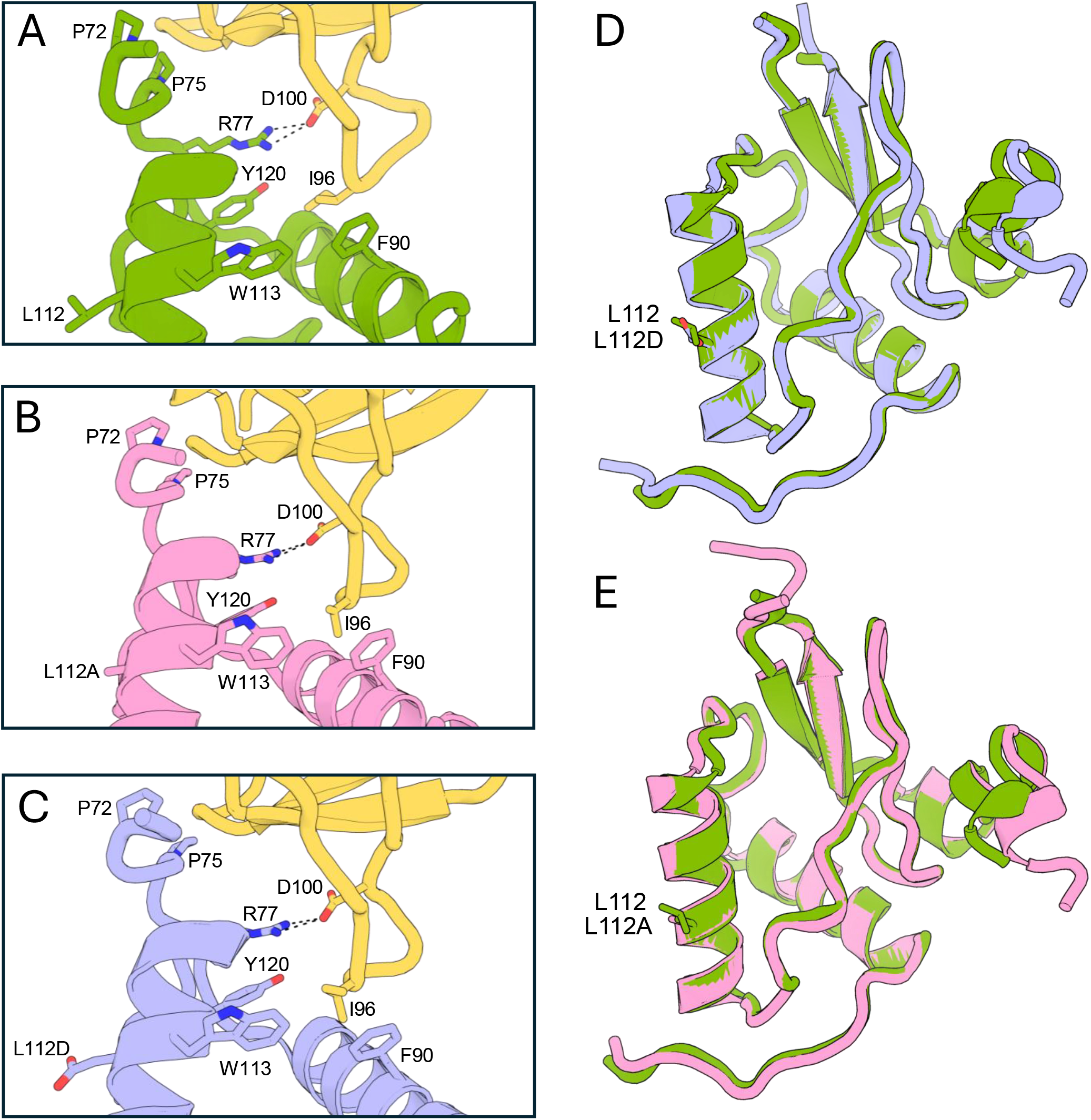
Nef core fold and SH3 interface are maintained in Nef mutant structures. Close-up view of SH3 domain (yellow) interfaces with wild-type Nef (**A**), Nef-L112D (**B**) and Nef-L112A (**C**). The Nef PxxPxR motif forms a polyproline type II helix that engages the SH3 domain. This interaction is stabilized by a salt bridge formed between Arg77 of Nef and Asp100 in the RT loop of SH3. Ile96 of the RT loop of SH3 engages in a small hydrophobic pocket in the Nef core region formed by Phe90, Trp113 and Tyr120. Structural alignment of wild-type Nef core (green) with Nef-L112D (**D**; periwinkle) and Nef-L112A (**E**; pink) revealed RMSD values of 0.79 and 0.82, respectively, across backbone α-carbons.

Alignments of the Nef-L112A and Nef-L112D structures with wild-type Nef were performed with US-align (24) to gauge the similarity of the core folds. Neither mutation affected the global fold of the Nef core region: the structures of Nef-L112D (Figure 3D) and Nef-L112A (Figure 3E) are virtually identical to that of wild-type, with overall root mean square deviations (RMSD) of just 0.82 and 0.79 Å, respectively, for the αC-backbones. These structures demonstrate that the L112A and L112D mutations abolish Nef dimerization while preserving the global fold of the Nef core and the Nef:SH3 interface. Taken together, they support the broader conclusion that the pleiotropic biological effects of Nef dimer interface mutations are due to the dimerization defect rather than perturbation of Nef:SH3 interaction.

### SplitFAST Complementation Assay Reports Nef Homodimer Formation in Solution

Previous studies have reported the discovery and development of small molecules that bind to Nef (7). One series of compounds, based on a hydroxypyrazole core, inhibits multiple Nef functions including enhancement of viral infectivity and replication as well as kinase activation and MHC-I downregulation (16–18). Table 3 summarizes the structures and activities of several compounds in this class, with analog FC-8052 showing the most promise based on the aggregation of all in vitro and cellular activity. The effects of these compounds on diverse Nef functions are reminiscent of those observed with the dimerization interface mutations, suggesting that they may also interfere with Nef homodimer formation as a mechanism of action. To explore this possibility, we developed an assay to monitor Nef homodimerization in solution. This assay is based on a small protein tag (fluorescence-activating and absorption-shifting tag, FAST; 13.6 kDa) (20) that produces a strong fluorescent signal in the presence of various rhodanine fluorogens. The FAST coding sequence was split into two non-fluorescent fragments (nFAST and cFAST) and fused to the C-terminal coding region of full-length Nef (Figure 4A). We tested cFAST tags of 9 and 11 amino acids in length (cFAST9 and cFAST11; Figure 4A) because previous studies have shown that the length of the cFAST tag impacts its intrinsic affinity for nFAST and self-complementation (21). The Nef-nFAST and Nef-cFAST fusion proteins were expressed in bacteria and purified to homogeneity. Complementary fusion protein pairs were then combined over a range of concentrations in equimolar ratios in the presence of the fluorogen, 4-hydroxy-3-methylbenzylidene-rhodanine (HMBR; assay principle illustrated in Figure 4B). Dimerization of Nef promoted complementation of the FAST fragments in both cases, resulting in HMBR binding and fluorescence, which increased as a function of the Nef-FAST fusion protein concentration (Figure 4C).

**Table 3.**
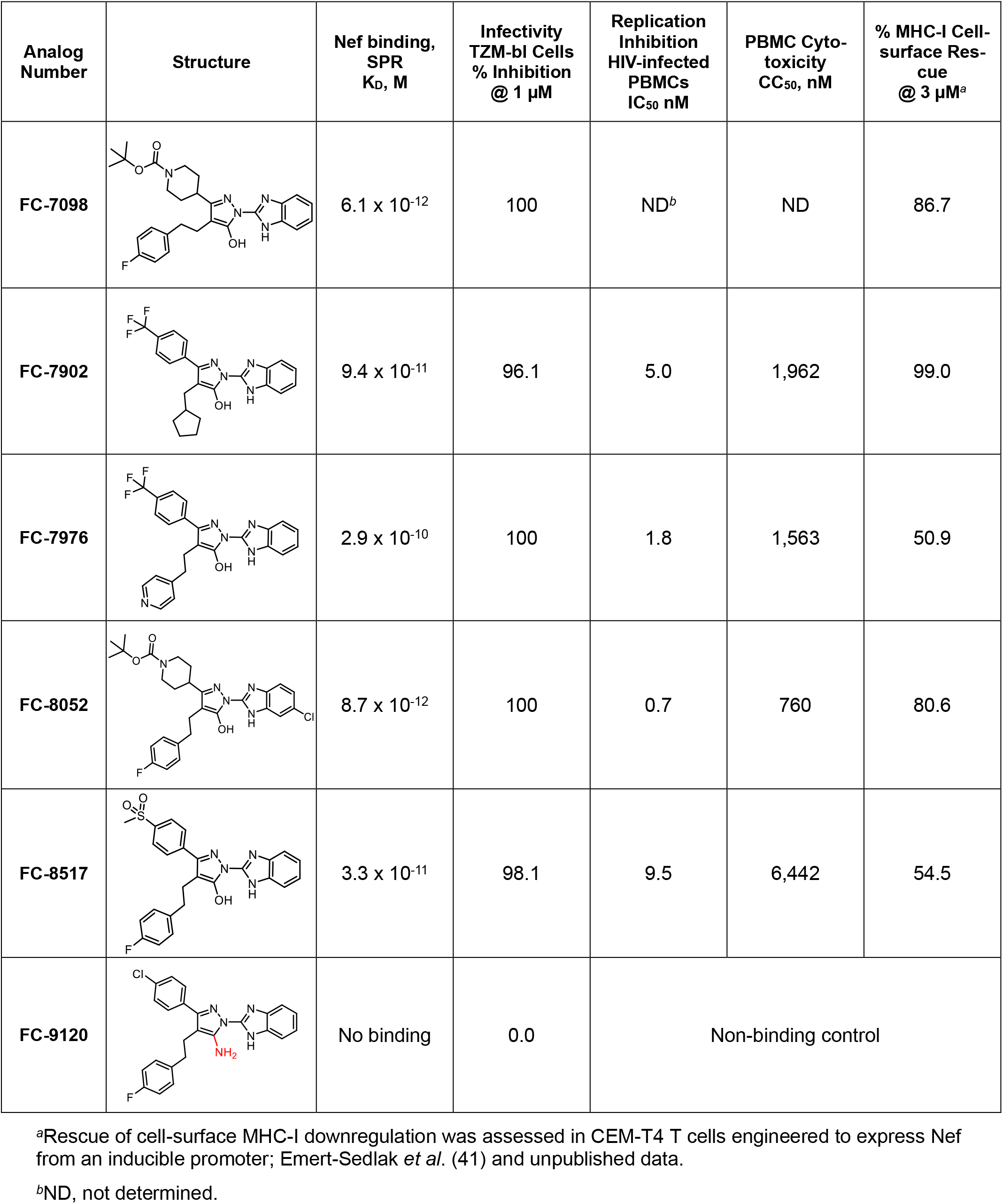
Properties of Nef Inhibitors used in this study. Synthesis and further characterization of these inhibitors are reported in Shi, *et al*. (16). *^a^*Rescue of cell-surface MHC-I downregulation was assessed in CEM-T4 T cells engineered to express Nef from an inducible promoter; Emert-Sedlak *et al*. (41) and unpublished data. *^b^*ND, not determined.

| Analog Number | Structure | Nef binding, SPR<br>$K_D$, M | Infectivity<br>TZM-bl Cells<br>% Inhibition<br>@ 1 μM | Replication<br>Inhibition<br>HIV-infected<br>PBMCs<br>$IC_{50}$ nM | PBMC Cyto-<br>toxicity<br>$CC_{50}$, nM | % MHC-I Cell-surface Res-<br>cue<br>@ 3 μM<sup>a</sup> |
| --- | --- | --- | --- | --- | --- | --- |
| FC-7098 | | $6.1 \times 10^{-12}$ | 100 | ND<sup>b</sup> | ND | 86.7 |
| FC-7902 | | $9.4 \times 10^{-11}$ | 96.1 | 5.0 | 1,962 | 99.0 |
| FC-7976 | | $2.9 \times 10^{-10}$ | 100 | 1.8 | 1,563 | 50.9 |
| FC-8052 | | $8.7 \times 10^{-12}$ | 100 | 0.7 | 760 | 80.6 |
| FC-8517 | | $3.3 \times 10^{-11}$ | 98.1 | 9.5 | 6,442 | 54.5 |
| FC-9120 |  | No binding | 0.0 | Non-binding control |  |  |

<sup>b</sup>ND, not determined.

**Figure 4.**
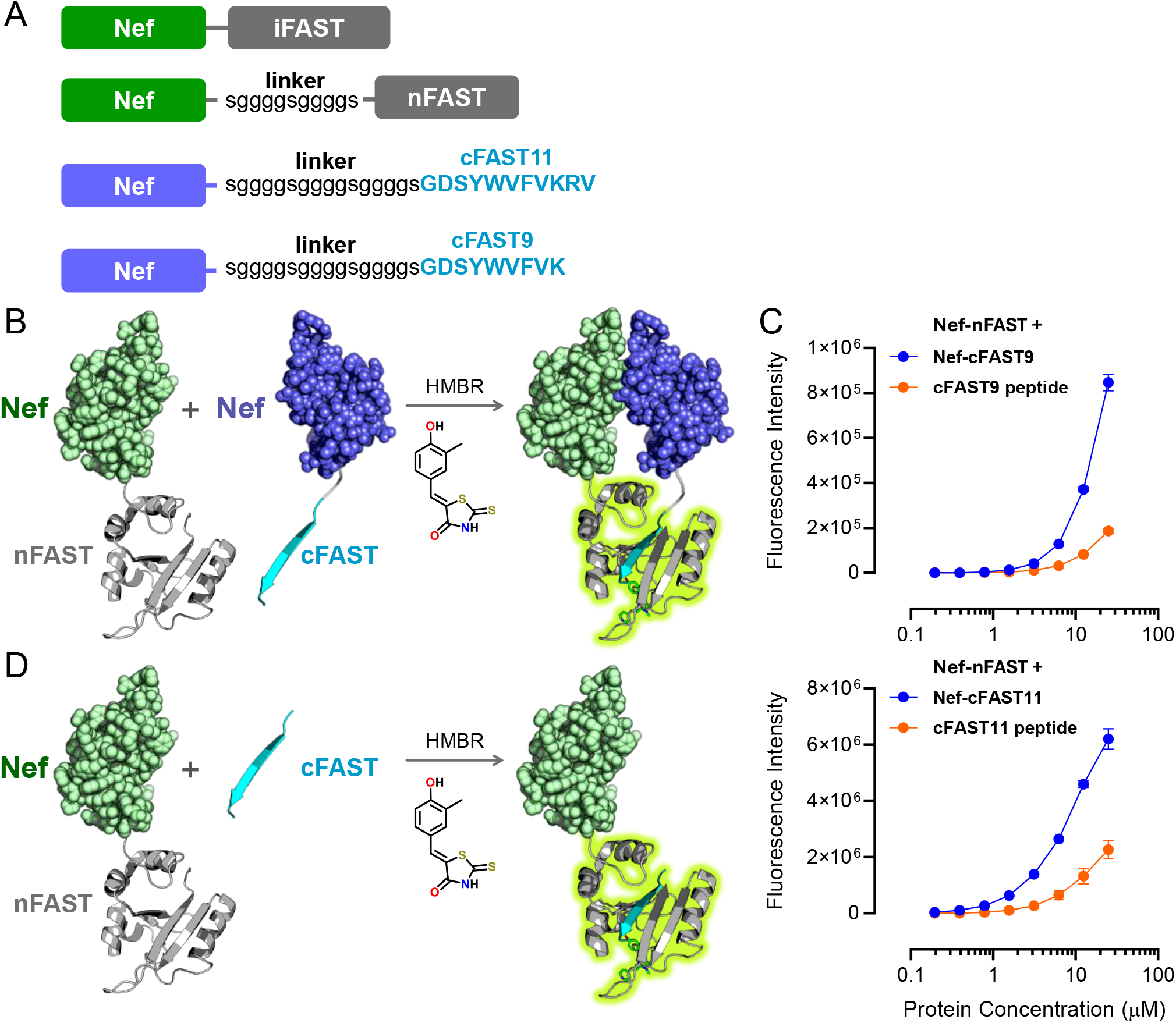
SplitFAST complementation assay for Nef homodimer formation in solution. **A.** Constructs used in a reversible split fluorescent reporter assay (splitFAST) to assess Nef homodimer formation. **B.** SplitFAST assay principle. Non-fluorescent nFAST and cFAST fragments are fused to full length Nef and incubated in the presence of the fluorogen, HMBR. Nef homodimer formation juxtaposes the FAST fragments, promoting complementation and fluorescence upon HMBR binding. **C.** Nef dimer formation was evaluated with Nef-nFAST and Nef-cFAST9 (*top panel*) or Nef-cFAST11 (*bottom panel*) fragments along with the corresponding cFAST9 and cFAST11 peptides for self-complementation controls. Nef-dependent complementation data points are shown in blue with self-complementation in orange. All data points were measured in quadruplicate and are shown as the mean value ± SE. **D.** Illustration of self-complementation control in which Nef-nFAST is mixed with the corresponding cFAST peptide without Nef.

To control FAST protein self-complementation in the absence of Nef dimerization, additional assays were conducted that combined Nef-nFAST with synthetic peptides consisting of only the cFAST9 and cFAST11 sequences (self-complementation illustrated in Figure 4D). Both cFAST peptides showed significantly reduced fluorescence when combined with Nef-nFAST compared to complementation with the corresponding Nef-cFAST fusion protein (Figure 4C), supporting Nef-dependent complementation. While Nef-cFAST9 generated a smaller complementation signal than Nef-cFAST11 when combined with Nef-nFAST, self-complementation with the cFAST11 peptide was proportionally higher, resulting in a smaller signal window for inhibitor assays. For this reason, subsequent experiments used Nef-cFAST9 constructs.

### Hydroxypyrazole Inhibitors Disrupt Nef Homodimer Formation

Using the splitFAST assay, we evaluated the effects of previously characterized hydroxypyrazole Nef inhibitors (Table 3) on Nef homodimer formation. Inhibitors FC-7902, FC-8517, FC-8052, and FC-7976 caused a significant concentration-dependent reduction in fluorescence intensity, indicating impaired Nef dimer formation (Figure 5A). Conversely, FC-9120, an analog with an aminopyrazole in place of the hydroxypyrazole core (Table 3), showed no effect on fluorescence complementation, consistent with previous studies demonstrating its lack of Nef binding and antiretroviral activity (Figure 5B).

**Figure 5.**
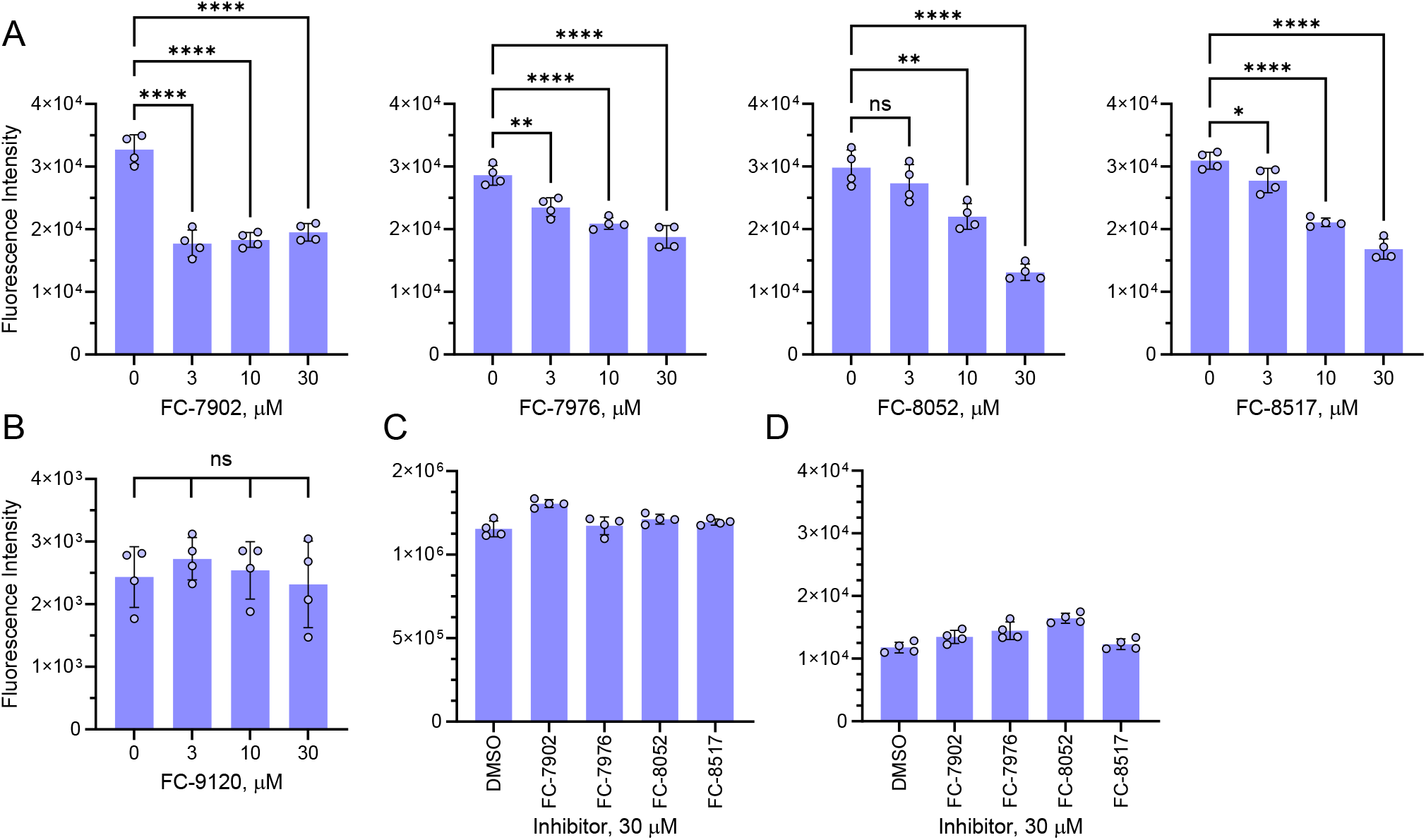
Small molecule inhibitors suppress Nef homodimer formation. **A.** Nef homodimer formation was assayed in the presence of inhibitors FC-7902, FC-7976, FC-8052, and FC-8517 at the concentrations shown. **B.** Result with non-binding control analog, FC-9120. **C.** Nef-iFAST, in which Nef is fused to the intact FAST protein (no complementation) was evaluated in the presence of the inhibitors to rule out fluorescence quenching. **D.** Each inhibitor was tested in the presence of Nef-nFAST and the cFAST9 peptide to rule out effects on self-complementation. In all panels, bar heights indicate the mean value of at least four replicates ± SE. Significant differences from the untreated controls were assessed by one-way ANOVA; *, p < 0.05; **, p < 0.01; ****, p < 0.0001; ns, not significant.

Additional control experiments support the conclusion that the reduced fluorescence observed in the presence of the inhibitors is due to Nef homodimer disruption. Full-length Nef fused to an intact FAST tag (no complementation) maintained a consistent fluorescent signal in the presence of all inhibitors at 30 μM, the highest concentration tested. This control rules out the possibility of fluorescence quenching or interference with fluorogen binding to the FAST tag (Figure 5C). Additionally, each inhibitor was tested against the fluorescence observed in the presence of Nef-nFAST and the unfused cFAST9 peptide. No significant changes were observed, confirming that the compounds do not impact self-complementation (Figure 5D). In addition, this experiment shows that the residual fluorescence observed in the presence of high inhibitor concentrations (Figure 5A) is most likely due to self-complementation. Overall, these results provide direct evidence that hydroxypyrazole inhibitors block multiple Nef functions by interfering with homodimer formation.

### Co-crystallization of the wild-type Nef:SH3 complex with inhibitor prevents dimer formation

Results presented in the previous section support a mechanism for Nef inhibitor action that involves suppression of homodimer formation, a key aspect of many Nef functions. To extend this result, we co-crystallized the wild-type Nef:SH3 complex in the presence of the hydroxypyrazole Nef inhibitor analog FC-7098, which is identical in structure to FC-8052 except for the chlorine atom on the benzimidazole moiety (Table 3). Unlike the 2:2 apo complex (Figure 1), wild-type Nef:SH3 crystallized as a 1:1 complex in the presence of the small molecule (Figure 6A) with no evidence of the Nef homodimer interface. The presence of the small molecule did not affect the interaction of Nef with SH3 ─ the Nef polyproline helix still binds the SH3 surface and Nef Arg77 forms the salt bridge with Asp100 in the SH3 domain RT loop in same manner observed in the wild-type Nef structure (Figure 6B). Structural alignment of the Nef core observed in the presence of FC-7098 with that of the apo core shows that the inhibitor did not influence the overall Nef fold (RMSD = 0.89 Å over the αC-backbone; Figure 6C). Despite the impact of FC-7098 on the quaternary structure of the Nef:SH3 complex, no electron density was observed for the inhibitor itself. Regardless, this outcome provides structural support for a mechanism of action by which hydroxypyrazoles interfere with Nef homodimer formation.

**Figure 6.**
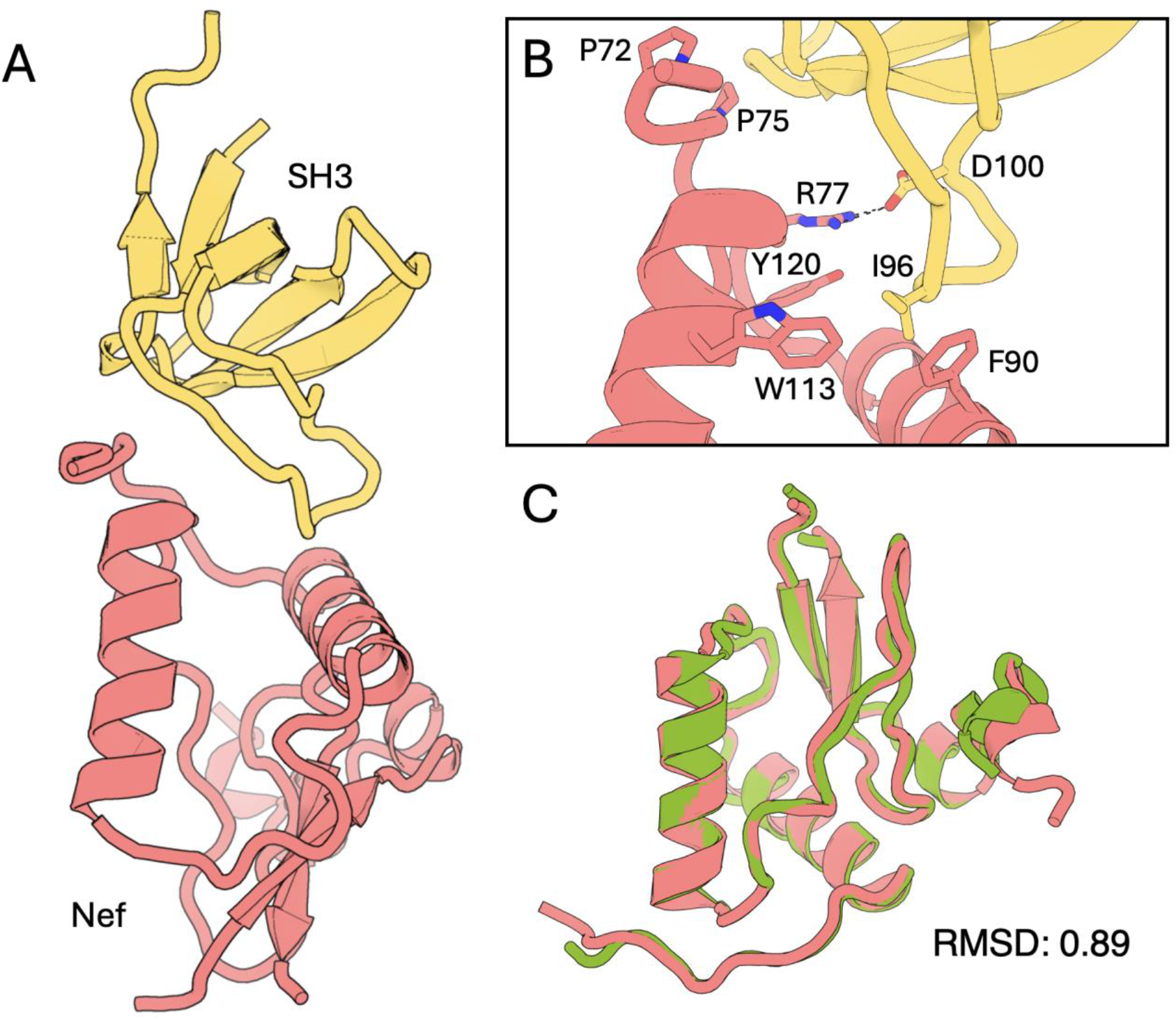
Co-crystallization of the Nef:SH3 complex with Nef inhibitor FC-7098 prevents dimer formation. The purified complex of the Nef core region with the Hck SH3 domain was co-crystallized with the hydroxypyrazole inhibitor FC-7098 (see Table 3 for structure). **A.** The overall complex crystallized with 1:1 Nef to SH3 stoichiometry. **B.** The presence of the inhibitor in the complex did not affect the Nef:SH3 interface, involving contacts formed by the Nef PPII helix (Pro72, Pro75, Arg77) and hydrophobic pocket (Phe90, Trp113, Tyr120) with the SH3 domain binding grooves and RT loop side chains Ile96 and Asp100. **C.** The Nef core from the FC-7098 co-crystallization structure (salmon) was aligned with the Nef core from the apo structure shown in Figure 1 (PDB: 1EFN; chain B, green). The alignment showed little difference, with an RMSD of 0.89 Å across backbone α-carbons. No electron density was observed for the inhibitor, which may reflect its lower affinity for monomeric Nef (see text for details).

The crystallography result, combined with the SplitFAST data, supports a model in which the inhibitors initially recognize the Nef dimer, followed by dimer disruption and inhibitor release. The model predicts that the inhibitors show enhanced affinity for the dimer vs. the monomeric form. To test this idea computationally, we performed unbiased docking of each of the active compounds shown in Table 3 against the entire dimeric structure of Nef present in PDB 1EFN with the SH3 domains removed. All five analogs docked to a pocket formed by the Nef dimer structure (Figures 7A and B). This pocket is adjacent to the helical dimer interface shown in Figure 1. The same docking routine was then repeated with each analog using a single monomer from the same crystal structure. In this case, the top scoring inhibitor poses mapped to multiple different sites, including the dimer interface itself as well as to a shallow groove on the opposite side of the Nef molecule. To evaluate the differences in predicted binding affinity, we compared the binding scores for the top inhibitor poses against the dimer vs. the monomer. The average of the predicted binding scores for the top poses with the dimer is -10.14 kcal/mol, while the docking scores with the monomer were significantly weaker with an average binding score of -8.04 kcal/mol (Figure 7C). This observation provides independent support for the idea that Nef inhibitors in this class initially recognize the dimer as a first step in a cycle leading to dimer disruption as shown by SplitFAST assay.

**Figure 7.**
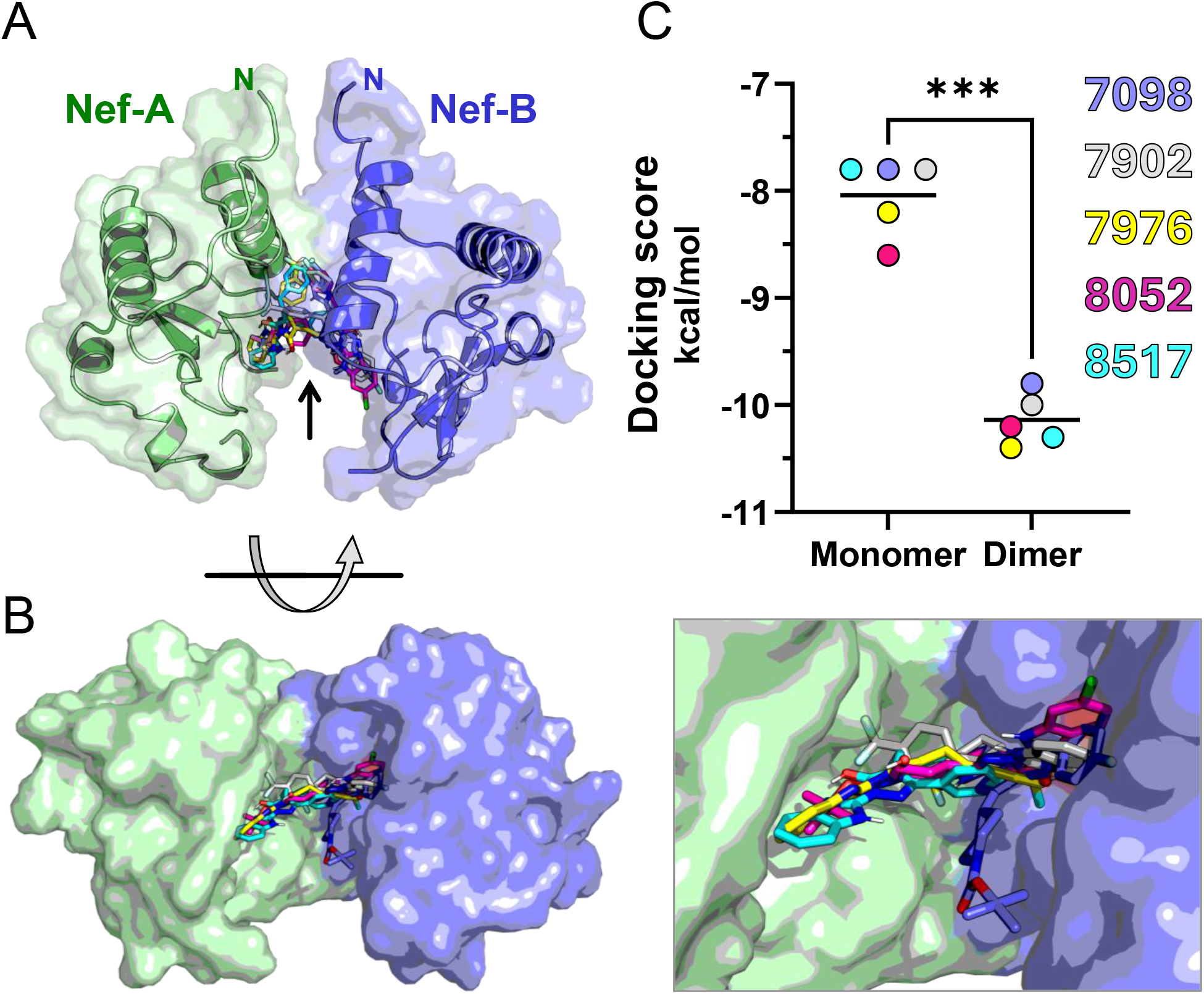
Unbiased docking supports preferential hydroxypyrazole Nef inhibitor recognition of the dimeric state. **A)** The five active Nef inhibitors shown in Table 3 were docked against the Nef dimer present in PDB 1EFN using MzDock in blind mode (see Materials and Methods). All five analogs docked to the same site which is located adjacent to the dimer interface (*arrow*). The Nef monomers are colored in green and blue, respectively, and the N-termini are indicated. **B)** The model in panel B is rotated 90° to show the binding pocket in the overall dimer (*left*) with a close-up view on the right. **C)** Docking was repeated with a single monomer from the same crystal structure, and the scores from the top poses with the monomer were compared with the dimer scores by paired t-test (***, p = 0.0001).

## Discussion

Our study sheds new light on the structural and mechanistic roles that homodimer formation plays in the function of Nef. Mutations that disrupt the dimer interface do not alter the global fold of the Nef core or its interaction with the Hck SH3 domain, supporting the conclusion that the phenotypic effects of these mutations can be attributed solely to the loss of the dimer. Consistent with these structural observations, previous work has shown that the same mutations in full-length Nef interfere with kinase activation and viral replication, most likely through a dominant-negative mechanism (10). Using a split fluorescent complementation assay, we were able to demonstrate homodimerization of full-length purified Nef in solution and show that hydroxypyrazole Nef inhibitors interfere with homodimerization as a likely mechanism of action in cells. Altogether, these findings highlight the Nef dimer interface as an attractive target for continued antiretroviral drug development targeting Nef.

The effects of hydroxypyrazole inhibitors on Nef function closely mirror the effects of dimerization-defective mutants, both of which block multiple Nef functions and lead to reduced HIV-1 infectivity and replication (8). Co-crystallization of wild-type Nef:SH3 complex with the prototypical inhibitor FC-7098 revealed a 1:1 Nef:SH3 structure with no evidence of the Nef homodimer interface. Much like the crystal structure of the Nef-L112A:SH3 complex presented here and a previous structure of a Nef-L112D:SH3 complex, both of which are also 1:1 complexes, the fold of the Nef core remained intact and the canonical SH3 binding site on Nef was undisturbed. Despite these effects of the inhibitor on the Nef complex, we did not observe electron density for FC-7098 itself. One possible explanation is that initial binding of the inhibitor requires the Nef dimer, followed by dimer disruption and release of the small molecule. Unbiased computational docking studies presented in Figure 7 support this idea. All five inhibitors were predicted to bind to a common binding pocket present in the dimer but not in the monomer. In contrast, docking against a single monomer resulting in significantly lower binding scores, with an average difference of about 2 kcal/mol. This difference translates to at least a 30-fold reduction in binding affinity, which is consistent with release of the small molecule from the monomer once the dimeric form is disrupted. Suppression of Nef homodimer formation in the splitFAST assay paired with structural insights from co-crystallization with FC-7098 as well as the docking results support a mechanism by which hydroxypyrazole Nef inhibitors interfere with dimer formation without affecting the core fold of Nef.

Previous structural biology studies have demonstrated that residues present in the Nef homodimer interface also form direct contact points for other host proteins, suggesting that this region of Nef serves as a multifunctional hub controlling dimer formation and effector protein recruitment. One important example involves the cellular machinery by which Nef drives CD4 downregulation from the cell surface. This process requires Nef-mediated assembly of a complex involving the AP-2 endocytic adaptor protein and the cytoplasmic C-terminal tail of CD4 (2). In a crystal structure of the AP-2 α/σ2 subunit hemicomplex with a Nef-CD4 fusion protein (PDB: 6URI) (25), Nef dimer interface residues, including Leu112, make direct contacts with CD4 and the homodimer is not present. Inhibitors or mutations targeting this region may impair Nef-mediated CD4 downregulation by directly interfering with complex assembly. The dual roles that Leu112 and nearby residues play in Nef function highlights their versatility as a scaffold for both homodimer formation and as a surface for stabilizing effector protein binding. Whether or not Nef homodimer formation represents a necessary intermediate in the pathway leading to AP-2 mediated downregulation of CD4 will require further investigation.

In contrast to the structure of Nef in complex with AP-2 and CD4, Nef dimerization may play an indirect role in the downregulation of MHC-I, which is mediated by the AP-1 endocytic adaptor (2). Alignment of the structures of one subunit of the Nef homodimer with a structure of a Nef-MHC-I fusion protein complexed with the AP-1 µ1 subunit shows that residues involved in the Nef homodimer interface do not interact directly with MHC-I or AP-1 (8). However, Asp123 of Nef plays a key role, forming part of an electrostatic network with Arg393 from AP-1 µ1 which in turn contacts Asp327 from the MHC-I tail. Based on the dimeric crystal structure of Nef in complex with the Hck SH3-SH2 region, interaction of the Nef homodimer with Hck may reorient Nef Asp123 from a buried position in the core to a solvent-exposed position compatible with MHC-I and AP-1 interaction. Structural alignment of one Nef monomer present in the Hck SH3-SH2 complex (PDB: 4U5W) (14) with Nef in the MHC-I/AP-1 complex (PDB: 4EN2) (26) revealed that Nef Asp123 adopts a nearly identical surface-exposed pose in both structures (27). This observation suggests that interaction of Nef with Hck or other Src-family members at the cell membrane not only leads to kinase activation via a dimerization-dependent mechanism but may also induce a conformation of the Nef core favorable for subsequent recruitment of MHC-I and AP-I for downregulation. Additional support for this idea comes from older work demonstrating a central role for Src-family kinase activation in one pathway leading to MHC-I downregulation (28, 29). Small molecule inhibitors of Nef that reverse Nef-mediated downregulation of MHC-I may therefore disrupt the initial kinase activation step in this pathway which is dependent on the homodimer.

## Materials and Methods

### Expression and Purification of Nef complexes with Hck SH3 and SH3-SH2 proteins

Nucleotide sequences encoding the core region of HIV-1 Nef (SF2 variant; residues 55-209 based on the Nef:SH3 complex crystal structure, PDB: 1EFN) and the SH3 (residues 56-119) or SH3-SH2 (residues 51-221) domains of the human p59 form of Hck (numbering as per UniProt ID: P08631-2) were amplified by PCR and subcloned into pET-based expression plasmids (14, 27). The SH3 and SH3-SH2 proteins include a C-terminal His tag to allow capture of the Nef-bound complexes. *E. coli* strain Rosetta2 (DE3) pLysS (Millipore Sigma) was transformed with the expression plasmid for the Nef core while *E. coli* strain BL21 Star (DE3) (ThermoFisher Scientific) was transformed with the Hck SH3 or SH3-SH2 tandem domain expression plasmids. For each construct, a single colony was picked and cultured overnight in 100 mL LB medium containing the appropriate antibiotic at 37 °C with shaking. Overnight cultures were used to inoculate 1 liter of LB medium supplemented with antibiotic and cells were grown at 37 °C to an OD_600_ of 0.6. The temperature was reduced to 18 °C for 30 min and protein expression was induced by adding IPTG to a final concentration of 0.5 mM. Cultures were shaken at 18 °C for 18 h. Cell pellets were harvested by centrifugation at 6,600 g for 15 min, flash-frozen, and stored at -80 °C.

To purify the Nef:SH3 and Nef:SH3-SH2 complexes, cell pellets were thawed on ice and resuspended in 50 mL of Ni-IMAC (nickel-immobilized metal affinity chromatography) binding buffer (25 mM Tris-HCl, pH 8.3, 0.3 M NaCl, 20 mM imidazole, 10% (v/v) glycerol, and 2 mM 2-mercaptoethanol). Nef:SH3 and Nef:SH3-SH2 cell suspensions were mixed, combined with one cOmplete protease inhibitor tablet (Roche), and passed through a microfluidizer (Microfluidics) 10 times at 4 °C. The cell lysates were incubated at 4 °C with gentle rocking for 1 h to promote complex formation followed by centrifugation at 100,000 g for 1 h at 4 °C. The clarified lysate was loaded onto a 5 mL His-Trap HP column (Cytiva) at 2.0 mL/min pre-equilibrated with Ni-IMAC binding buffer. Bound protein was eluted with a linear gradient of 20 to 500 mM imidazole in Ni-IMAC elution buffer (binding buffer plus 500 mM imidazole). Fractions containing both Nef and Hck proteins were identified by SDS-PAGE, pooled and concentrated to 1 mL with an Amicon 50 mL stirred-cell concentrator (10 kDa molecular weight cutoff; Millipore Sigma). The protein complexes were buffer-exchanged twice with gel filtration buffer (25 mM Tris-HCl, pH 8.0, 200 mM NaCl, 10% glycerol, 2 mM TCEP) and centrifuged at 27,000 g for 10 min at 4 °C. Supernatants containing the soluble protein complexes were loaded onto a Hi-Load 16/60 Superdex 75 gel filtration column (Cytiva) equilibrated with gel filtration buffer and eluted at a flow rate of 0.5 mL/min. Fractions containing the complexes were pooled, concentrated, flash-frozen in liquid nitrogen, and stored at −80 °C.

### Size exclusion chromatography-multi-angle light scattering (SEC-MALS)

SEC-MALS was performed using a Wyatt Technology instrument consisting of a DAWN multi-angle light scattering detector, a DLS-DAWN dynamic light scattering module for determination of hydrodynamic radius, and an Optilab refractometer. Purified Nef:SH3 and Nef:SH3-SH2 complexes (50 µL of a 10 µg/mL solution) were loaded onto a Superdex 75 Increase 10/300 GL column pre-equilibrated with SEC-MALS buffer (20 mM Tris-HCl, 0.15 M NaCl, 2 mM TCEP, 5% glycerol, pH 8.3). Data were recorded for 50 min at a flow rate of 0.5 mL/min at room temperature. Data collection and SEC-MALS analysis were performed with ASTRA 7.3.2 software (Wyatt Technology).

### Crystallography and X-ray data collection

For the Nef-L112A:SH3 complex, crystals were grown by hanging-drop vapor diffusion at room temperature by mixing the protein complex (9.7 mg/mL; 376 μM) in a 1:1 ratio with 0.1 M magnesium acetate, 0.1 M sodium acetate pH 4.6, and 25% (v/v) PEG 400. Large, single crystals of the complex grew in 14 days. Crystals were cryoprotected in 50 mM magnesium acetate, 50 mM sodium acetate pH 4.6, 12.5% (v/v) PEG 400, and 50% (v/v) xylitol prior to flash freezing in liquid nitrogen. X-ray diffraction data were collected at GM/CA-XSD beamline 23-ID-D at the Advanced Photon Source, Argonne National Laboratory. Data indexing, integration, and scaling were conducted using XDS (30). Diffraction data are consistent with the tetragonal P 4_1_ 2_1_ 2 space group. Solvent content analysis suggests 58.5% solvent and a Matthews coefficient of 2.96 Å^3^/Da.

Crystals of the wild-type Nef:SH3 complex in the presence of inhibitor FC-7098 were grown by sitting-drop vapor diffusion at room temperature. Recombinant purified Nef:SH3 complex (16.3 mg/mL; 628 µM) was incubated with Nef inhibitor FC-7098 (726 µM; 3.63% DMSO) at a 1:1.1 molar ratio overnight at 4 °C. The Nef:SH3:FC-7098 mixture was centrifuged at 15,000 g at 4 °C for 5 min to pellet insoluble material. Co-crystallization was carried out by mixing the Nef:SH3:FC-7098 sample in a 1:1 ratio with the crystallization solution (0.162 M sodium nitrate, 16.2% (w/v) PEG 3350, 2.7% (v/v) 2-methyl-2,4-pentanediol (MPD), 2.5% v/v ethylene glycol). Crystals formed in eleven days and were cryoprotected by a rapid immersion in 0.162 M sodium nitrate, 16.2% (w/v) PEG 3350, 25% (v/v) MPD, 2.5% v/v ethylene glycol followed by flash cooling in liquid nitrogen. X-ray diffraction data were collected using the University of Pittsburgh X-ray facility home source. Crystals diffracted X-rays to 2.34 Å and the data were indexed, integrated and scaled using XDS. Diffraction data are consistent with the tetragonal P4_1_2_1_2 space group. Solvent content analysis suggests 54.2% solvent and a Matthews coefficient of 2.69 Å^3^/Da, which is consistent with one Nef:SH3 complex in the asymmetric unit.

### Structure determination and refinement

For the Nef-L112A:SH3 complex, the structure factor data were analyzed using PHENIX Xtriage (31). Phasing and structure solution were conducted by molecular replacement with PHASER (32) using the structure coordinates of an individual Hck SH3 domain from PDB ID: 4U5W and of an HIV-1 Nef core from PDB ID: 1EFN. The top molecular replacement solution generated a singular Nef:Hck SH3 complex. After initial refinement, a preliminary model was generated with the Nef-L112A:SH3 sequence using the AlphaFold model prediction and Process Predicted Model tools in PHENIX (33). The processed model was used as a reference throughout refinement. Water molecules were added using phenix.refine after the penultimate cycle of refinement and model building. The model was built using Coot (34) and the final refined model was evaluated with MolProbity (35). Models of X-ray structures were created using PyMol (Schrödinger). No density was observed for the flexible internal loop of Nef (residues 156-180) which is consistent with previous Nef core crystal structures not bound to the AP2 adaptor protein complex.

The structure of the wild-type Nef:SH3 complex co-crystallized with FC-7098 was determined in a similar manner. However, the search models used for molecular replacement were the coordinates for the Nef-SF2 core and Hck SH3 domain from the Nef:Hck SH3-SH2 complex structure (PDB ID: 4U5W). One Nef:SH3 solution was found and refinement of this initial model was conducted with phenix.refine using rigid-body, cartesian simulated-annealing, individual coordinate, b-factor and occupancy refinement. Subsequent rounds of model building were conducted using Coot and refinement included TLS, individual coordinate, b-factor and occupancy refinement. Three ethylene glycol, one nitrate ion and thirty-one water molecules were modelled in the structure.

During structure refinement and improvement of the model, F_o_-F_c_ maps were analyzed to identify FC-7098 inhibitor-positive density but none was observed in the structure. Additional attempts to identify electron density of FC-7098 included refinement using data at lower resolution (3 Å) with improved I/σ followed by analysis of F_o_-F_c_ maps and calculating Polder maps to exclude bulk solvent around residue side-chains that make Nef dimer contacts. Ultimately no density was observed for the FC-7098 ligand.

### Bacterial expression constructs for the SplitFAST assay

All expression constructs were generated using Gibson cloning (Thermo Fisher). The coding sequence for full length HIV-1 Nef (SF2 variant) was cloned into the pSMT3 bacterial expression vector to introduce an N-terminal His6-small ubiquitin-like modifier (SUMO) tag. Various SplitFAST coding sequences were fused to the Nef C-terminus for expression of Nef-iFAST (complete iFAST sequence), as well as the Nef-nFAST, Nef-cFAST9 and Nef-cFAST11 fusion proteins. The amino acid sequences of iFAST and nFAST are based on the work of Tebo et al. (21) while the amino acid sequences of the cFAST peptides are presented in Figure 4. All expression constructs were verified by Sanger sequencing. Synthetic cFAST peptides used for control experiments were synthesized by the University of Pittsburgh Peptide Synthesis Core.

### Expression and Purification of Nef-FAST protein constructs

*E. coli* strain Rosetta 2 (DE3) pLysS (Millipore Sigma) was transformed with each Nef-FAST fusion protein expression vector. For each construct, a single colony was picked and cultured overnight in 100 mL LB medium at 37 °C in the presence of kanamycin. The overnight culture was then used to inoculate 1 L of LB medium supplemented with kanamycin. The cells were grown to an OD_600_ of 0.6 at 37 °C and the temperature was reduced to 18 °C for an additional 30 min with shaking. Protein expression was then induced by adding IPTG (0.5 mM final concentration) followed by overnight shaking at 18 °C. The cell pellets were collected by centrifugation at 6,600 g for 15 min, snap frozen and stored at -80 °C.

Each cell pellet was resuspended in 40 mL Ni-IMAC Buffer A (25 mM Tris-HCl, 0.5 M NaCl, 20 mM imidazole, 2 mM β-mercaptoethanol (BME), 10% glycerol, pH 8.3) and supplemented with one c0mplete protease inhibitor tablet (Roche). All remaining steps were carried out at 4 °C. Cells were lysed using a microfluidizer followed by ultracentrifugation at 100,000 g for 1 h to clarify the lysate. The clarified supernatant was loaded onto a 5.0 mL HisTrap HP column (Cytiva) pre-equilibrated with Ni-IMAC Buffer A. The column was then washed with 20 column volumes (CVs) of Ni-IMAC Buffer A. The His-tagged protein was eluted using a linear gradient combining Ni-IMAC Buffer A and Ni-MAC Buffer B (25 mM Tris-HCl, 0.5 M NaCl, 0.5 M imidazole, 2 mM BME, 10% glycerol, pH 8.3) over 32 CVs. Column fractions containing His6-SUMO-Nef-FAST proteins were identified by SDS-PAGE, pooled, and dialyzed overnight against Ni-IMAC dialysis buffer (25 mM Tris HCl, 0.5 M NaCl, 2 mM BME, 10% glycerol, pH 8.3) to remove the imidazole. Dialyzed protein was incubated with 50 µL of 2 mg/mL purified recombinant Ulp1 protease for 1 h and then loaded onto a 5.0 mL HisTrap HP column pre-equilibrated with Ni-IMAC Buffer A. The flow-through fraction containing the cleaved Nef-FAST protein was collected, while the His6-SUMO tag and His6-Ulp1 were retained by the column. The cleaved Nef-FAST protein was dialyzed against SEC Buffer (20 mM Tris-HCl, 0.15 M NaCl, 2 mM TCEP, 10% glycerol, pH 8.3) overnight at 4 °C and concentrated to a final volume of 5 mL using an Amicon stirred cell concentrator (Millipore Sigma) equipped with a cellulose membrane filter (3 kDa MWCO; Millipore Sigma) and Amicon Ultra-15 10 kDa MWCO centrifugal filters (Millipore Sigma). The concentrated protein was loaded onto a HiLoad 16/600 Superdex 75 pg preparative SEC column (Cytiva) pre-equilibrated with SEC buffer. Purified Nef-FAST protein fractions were identified by SDS-PAGE, pooled and concentrated to final concentrations of 32.0 mg/mL for Nef-iFAST, 12.0 mg/mL for Nef-nFAST, 7.7 mg/mL for Nef-cFAST9, and 7.1 mg/mL for Nef-cFAST11. Purified protein aliquots were flash-frozen in liquid nitrogen, and stored at -80 °C.

### Computational docking of inhibitors to Nef dimer and monomer

Docking was performed with the open-source software MzDock (36) which uses Smina (37), an enhanced version of AutoDock Vina (38), as the core docking engine. The Nef dimer crystal coordinates from PDB: 1EFN were prepared for docking by deleting the coordinates for the associated SH3 domains using PyMol (Schrödinger) (39). The monomeric Nef protein was then prepared by deleting the coordinates for one Nef subunit (chain D in 1EFN). The proteins were exported from PyMol as .CIF files which are compatible with the MzDock interface. Each of the Nef inhibitor analog structures shown in Table 3 were converted to SMILES strings using ChemDraw (Revvity). Docking was run in unbiased mode with the search box encompassing the entire Nef dimer or monomer. MzDock parameters were set to 9 modes per analog with exhaustiveness set to 32 using the MMFF94 force field for energy minimization (40). The predicted binding energies for the top scoring poses of the nine generated for each inhibitor with the monomer and the dimer were averaged and the statistical significance evaluated by pairwise Student’s t test (GraphPad Prism).

### SplitFAST assay for Nef dimerization and inhibitor action

All assays were performed in black 384-well microplates (Corning Cat. #3575) with final assay volumes of 20 µL/well. Initial titrations were performed by combining equimolar amounts of Nef-nFAST with either Nef-cFAST9 or Nef-cFAST11. Each protein pair was diluted to a combined concentration of 50 µM in Assay Buffer (20 mM Tris-HCl, 0.15 M NaCl, 2 mM TCEP, 10% glycerol, pH 8.3) and diluted in a series of 2-fold steps. Nef-nFAST was titrated with the corresponding cFAST9 and cFAST11 control peptides under identical conditions. Aliquots of each dilution (10 µL) were then combined with 10 µL of 10 µM HMBR prepared in Assay Buffer (TF-Lime; Twinkle Factory) in quadruplicate wells. Following incubation at room temperature for 1 h, fluorescence intensity was measured with an excitation wavelength of 480 nm and an emission wavelength of 541 nm on a Biotek Citation 5 plate reader.

Nef inhibitors were synthesized as described elsewhere (16). Each analog was prepared in 100% DMSO as a 10 mM stock and diluted to 100 µM using Assay Buffer. Inhibitor dilutions were combined with Nef-FAST protein pairs in Assay Buffer containing 1% DMSO in a 96-well plate at 2X the final desired concentration. Aliquots of each dilution were then combined with 10 µL of 10 µM HMBR in a 384-well microplate and fluorescence recorded as described above. All conditions were assayed in quadruplicate.

## Data availability

All data are described in the manuscript. The crystal coordinates for the structures of the Nef-L112A:SH3 and Nef:SH3 + FC-7098 complexes have been deposited in the Protein Data Bank and will be released upon publication (PDB IDs 9Y58 and 9Y9K, respectively).

## Funding

This work was supported by NIH grant AI152677 (to T.E.S). This research used resources of the Advanced Photon Source (APS) a U.S. Department of Energy (DOE) Office of Science User Facility operated for the DOE Office of Science by Argonne National Laboratory under Contract No. DE-AC02-06CH11357. The Eiger 16M detector at GM/CA-XSD was funded by NIH grant S10 OD012289GM/CA@APS is funded by the National Cancer Institute (ACB-12002) and the National Institute of General Medical Sciences (AGM-12006, P30GM138396). The content of this paper is solely the responsibility of the authors and does not necessarily represent the official views of the National Institutes of Health.

## Notes

The authors declare no competing financial interests.

## Notes

### Competing Interest Statement

The authors have declared no competing interest.

